# Occurrence and classification of T-shaped interactions between nucleobases in RNA structures

**DOI:** 10.1101/2022.10.22.513363

**Authors:** Zakir Ali, Teagan Kukhta, Ayush Jhunjhunwala, John F. Trant, Purshotam Sharma

## Abstract

Understanding the frequency and structural context of discrete noncovalent interactions between nucleotides is of pivotal significance in establishing the rules that govern RNA structure and dynamics. Although T-shaped contacts (*i.e*., perpendicular stacking contacts) between aromatic amino acids and nucleobases at the nucleic acid–protein interface have recently garnered attention, the analogous contacts within the nucleic acid structures have not been discussed. In this work, we have developed an automated method for identifying and unambiguously classifying T-shaped interactions between nucleobases. Using this method, we identified a total of 3262 instances of T-shaped (perpendicular stacking) contacts between two nucleobases in an array of RNA structures from an up-to-date dataset of high-resolution crystal structures deposited in the Protein Databank. These analyses add to the understanding of the physicochemical forces that are responsible for structure and dynamics of RNA.

## INTRODUCTION

Highly complex RNA 3D architectures are supported by an array of intramolecular forces (Holley et al. 1965; Tinoco Jr and Bustamante 1999), which include noncovalent interactions such as pairing (i.e., hydrogen bonding) and parallel π–π stacking of bases (Fig. 1 (Šponer et al. 1996; Leontis and Westhof 2001; Sharma et al. 2007; Sarver et al. 2008; Stombaugh et al. 2009; Havrila et al. 2013; Waleń et al. 2014; Jhunjhunwala et al. 2021)). Further, higher order hydrogen bonding (such as triplets and multiplets (Das et al. 2006; Abu Almakarem et al. 2012)) and stacking (e.g., base-intercalated, and base-wedge interactions (Baulin et al. 2020)) motifs determine RNA architecture. However, other interactions are also significant for RNA structure and function (Zirbel et al. 2009; Ulyanov and James 2010; Šponer et al. 2012). These include the interaction of metal ions with the phosphate groups, aiding RNA folding by minimizing the electrostatic repulsion (Agris 1996; Misra and Draper 1998; Draper 2004; Kolev et al. 2018), base–phosphate interactions (Zirbel et al. 2009), water-nucleobase stacking interactions (lone pair-π and OH-π interactions (Kalra et al. 2020)), ribose-nucleobase lone pair-π stacking interactions (Chawla et al. 2017), and T-shaped RNA inter-nucleobase interactions (Fig. 1). This last class, far less studied than the others, involves interactions between two perpendicularly interacting aromatic rings, in which one nucleobase skeleton provides the π face (denoted the ‘horizontal base’ in the subsequent text) and the other base an aryl group that interacts *via* its edge (denoted the ‘vertical base’, Fig. 1C).

**Figure 1.**
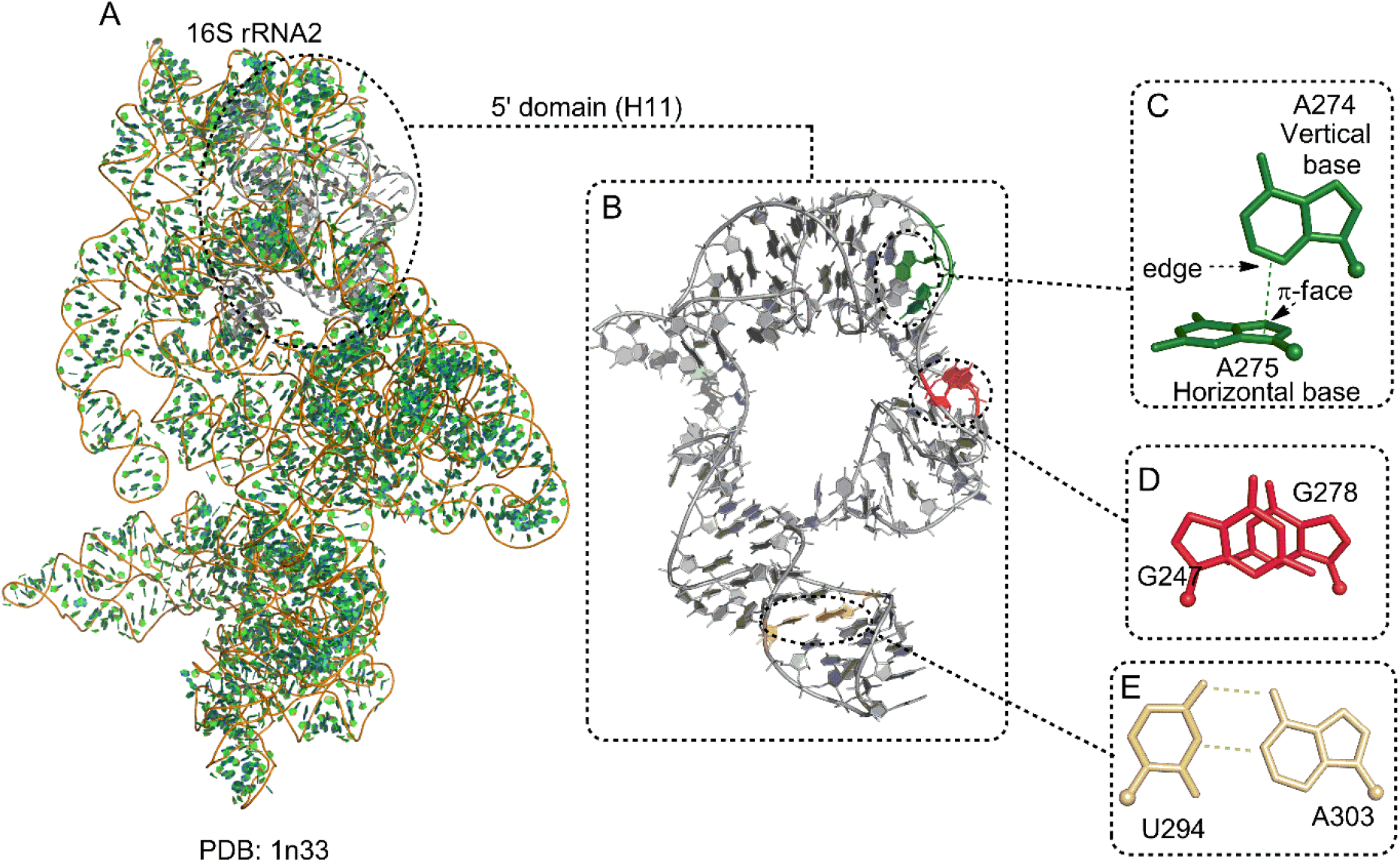
(A) Representation of the structure of 16S rRNA structure (PDB code: 1n33) belonging to the 30S ribosomal subunit of *Thermus thermophilus*, highlighting the nucleobase-specific noncovalent interactions in its (B) 5’ domain (H11). Examples of (C) T-shaped interactions, (D) base stacking and (E) base-pairing interactions are shown.

After their initial discovery in aromatic organic compounds (Sinnokrot and Sherrill 2004; DiStasio Jr et al. 2007; Geng et al. 2010), T-shaped interactions were identified in biomolecules, e.g., between two aromatic amino acids in proteins (McGaughey et al. 1998; Chelli et al. 2002), and between an aromatic amino acid and a nucleobase in nucleic acid–protein complexes (Rutledge and Wetmore 2008; Rutledge et al. 2009; Rutledge et al. 2010; Wilson et al. 2014; Wilson et al. 2016). For example, Wilson *et.al*. identified T-shaped interactions in a large crystal structure dataset of DNA-protein complexes and revealed their presence in the active sites of crucial enzymes (Wilson et al. 2014; Wilson et al. 2016). Further, Rutledge *et al* analysed T-shaped interactions between canonical nucleobases and aromatic amino acids using quantum mechanical calculations and revealed that their magnitude can be close to that of hydrogen bonds (Rutledge et al. 2009). The effects of methylation and the nature of monomer edge and face on T-shaped interactions were also analysed in nucleic acid–protein complexes (Rutledge and Wetmore 2008). However, no structural analysis exists on the T-shaped interactions between two nucleobases in RNA structures. As these are clearly critical in certain contexts, without this analysis, we cannot obtain a full picture of the energetics of the interactions within RNA complexes.

In this present work, we build on the existing RNA base pairing and parallel π–π stacking classification schemes to propose a simple and unambiguous geometry-based theoretical framework for the classification of T-shaped interactions in RNA (Leontis and Westhof 2001; Jhunjhunwala et al. 2021). We propose a five-stage hierarchy to classify them: **1)** identity of the interacting π face of the horizontal nucleobase; **2)** identity of the interacting edge of the vertical nucleobase; **3)** the mutual orientation of the glycosidic bonds of the two bases; **4)** the identity of the interacting ring(s) (five-membered, six-membered, or both) that constitute the horizontal base; and **5)** the relative position of the bases in the sequence (*consecutive, non-consecutive* and *inter RNA*, Fig. 2). Using this classification scheme in conjunction with the appropriately adapted existing approaches toward the detection of parallel π–π stacks (Jhunjhunwala et al. 2021) and T-shaped contacts (Wilson et al. 2014) in macromolecular 3D structures, we developed an algorithm for the automated identification and classification of examples of different T-shaped contacts in RNA crystal structures.

**Figure 2.**
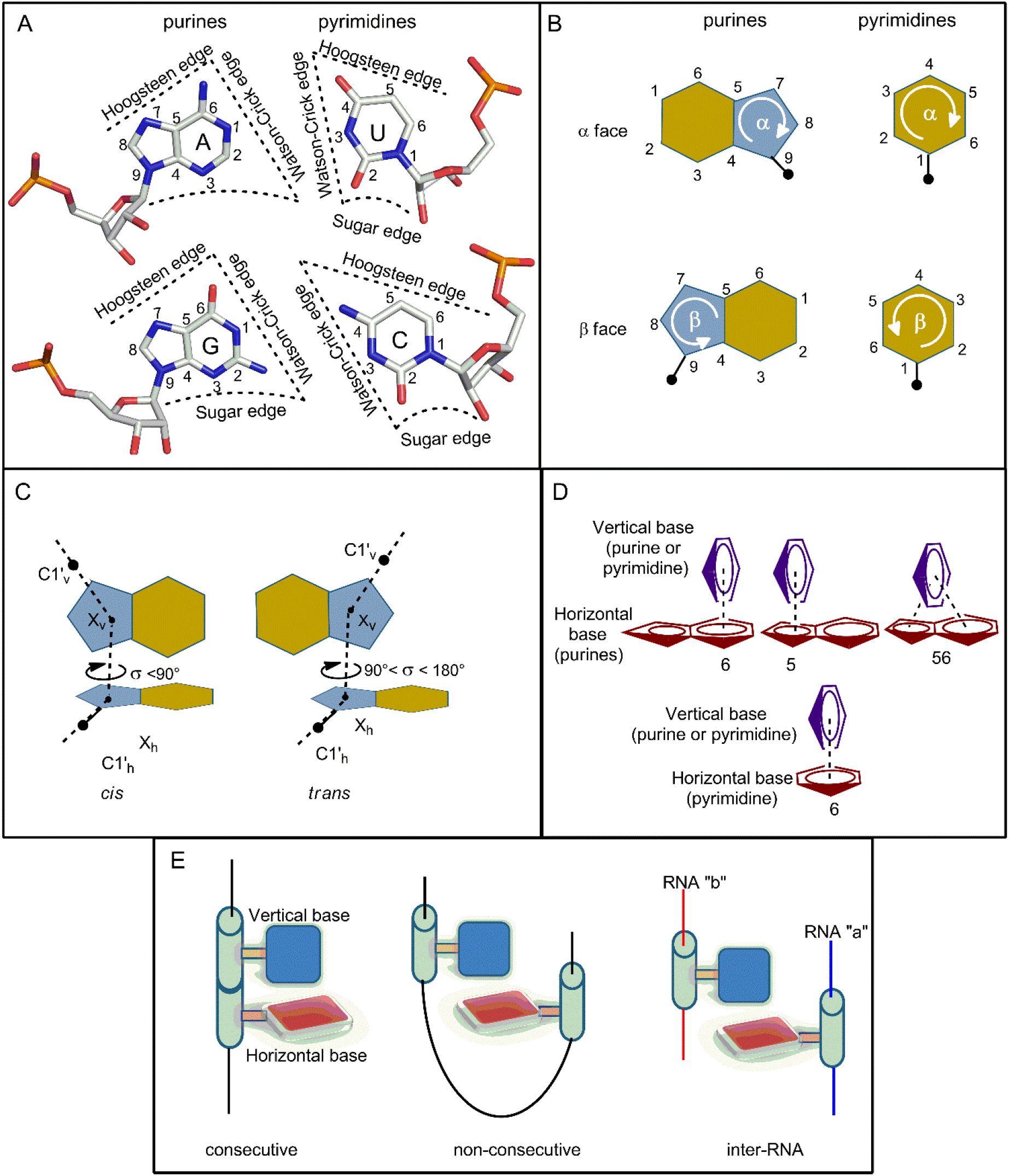
Representation of (A) edges, (B) faces, (C) relative glycosidic orientations, (D) identity of the participating horizontal ring, and (E) relative positioning in the sequence space for the nucleobases participating in a T-shaped interaction,

In this article, we describe the classification scheme and the algorithm. We then apply this scheme to analyse the occurrence and distribution T-shaped interactions in a non-redundant dataset of 2286 RNA structures with respect to the different structural parameters. We draw attention to the role that the T-shaped associations appear to play in connecting RNA secondary structural units in the 3D space. Our data reveals that these interactions play widely different roles in determining structure in different RNA types. This serves as a starting point for further multiscale investigations and/or simulations of RNA folding processes, and greater appreciation of the functional role of these interactions in biology.

## RESULTS AND DISCUSSION

### Classification of T-shaped interactions involving nucleobases

We searched for T-shaped interactions in an up-to-date dataset of RNA crystal structures. The detected T-shaped interactions were classified in several stages. In the first stage, the interactions were assigned to a basic geometric family based on the edge of the vertical base, the face of the horizontal base and the mutual glycosidic orientation of the interacting nucleotides (Fig. 2A-C). In the second stage, the geometries were assigned to subclasses based on the participation of the (five membered, six membered or both) rings that constitute a purine positioned horizontally in a T-shaped interaction (Fig. 2D). In the last stage, these interactions were further classified as *consecutive, non-consecutive*, or *inter-RNA* (Fig. 2E).

### Basic T-shaped Geometric Combinations

#### Combination of the interacting edge and face

A T-shaped contact in RNA involves the Watson-Crick (W) edge, Hoogsteen (H) edge or the Sugar (S) edge of the vertical base and the π-containing face (α or β (Rose et al. 1980)) of the horizontal base. For instance, if the ‘W’ edge of the vertical base interacts with the ‘α’ face of the horizontal base, the T-shaped interaction will be denoted as Wα or αW, depending on whether the horizontal or the vertical base occurs first in the RNA sequence. Assuming that the horizontal base occurs first in the sequence, each of the sixteen sequence combinations (i.e., four purine-purine (A-A, A-G, G-A and G-G), four purine-pyrimidine (A-C, A-U, G-C, G-U), four pyrimidine-purine (C-A, C-G, U-A and U-G), and four pyrimidine-pyrimidine (C-C, C-U, U-C, and U-U)) can have 6 distinct edge-face pairs: αW, αH, αS, βW, βH and βS, or 96 total possible combinations. *Relative glycosidic twist*. In analogy with the previously proposed RNA base pairing (Leontis and Westhof 2001) and stacking (Jhunjhunwala et al. 2021) classification schemes, T-shaped interactions can be divided in terms of the relative glycosidic torsion angle (σhv). Specifically, a T-shaped interaction can be annotated as *cis* if 90° ≥ σ_hv_ ≥ 0° and *trans* if 180° ≥ σ_hv_≥ 90° (Fig. 3). This leads to twelve basic T-shaped geometric families for each of the sixteen nucleobase sequence combinations (Table 1, *vide supra*).

**Figure 3.**
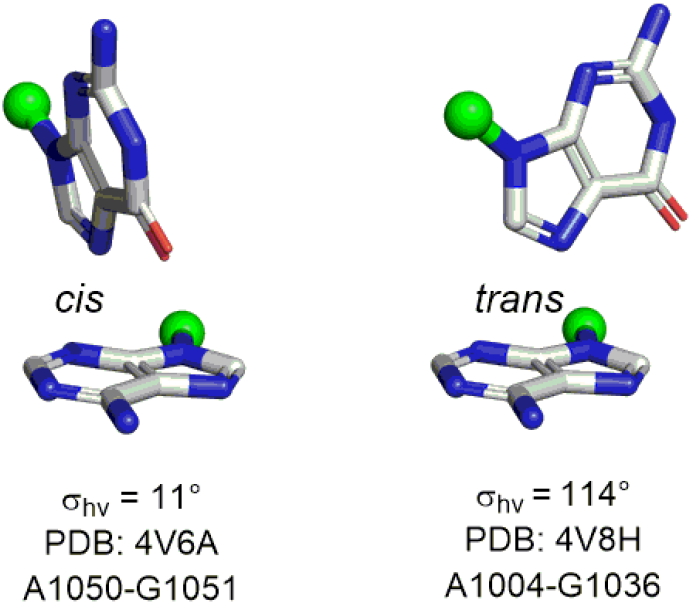
Examples of *cis* and *trans* T-shaped interactions. PDB code and interaction code (horizontal base-vertical base) is provided for each interaction.

**Table 1.**
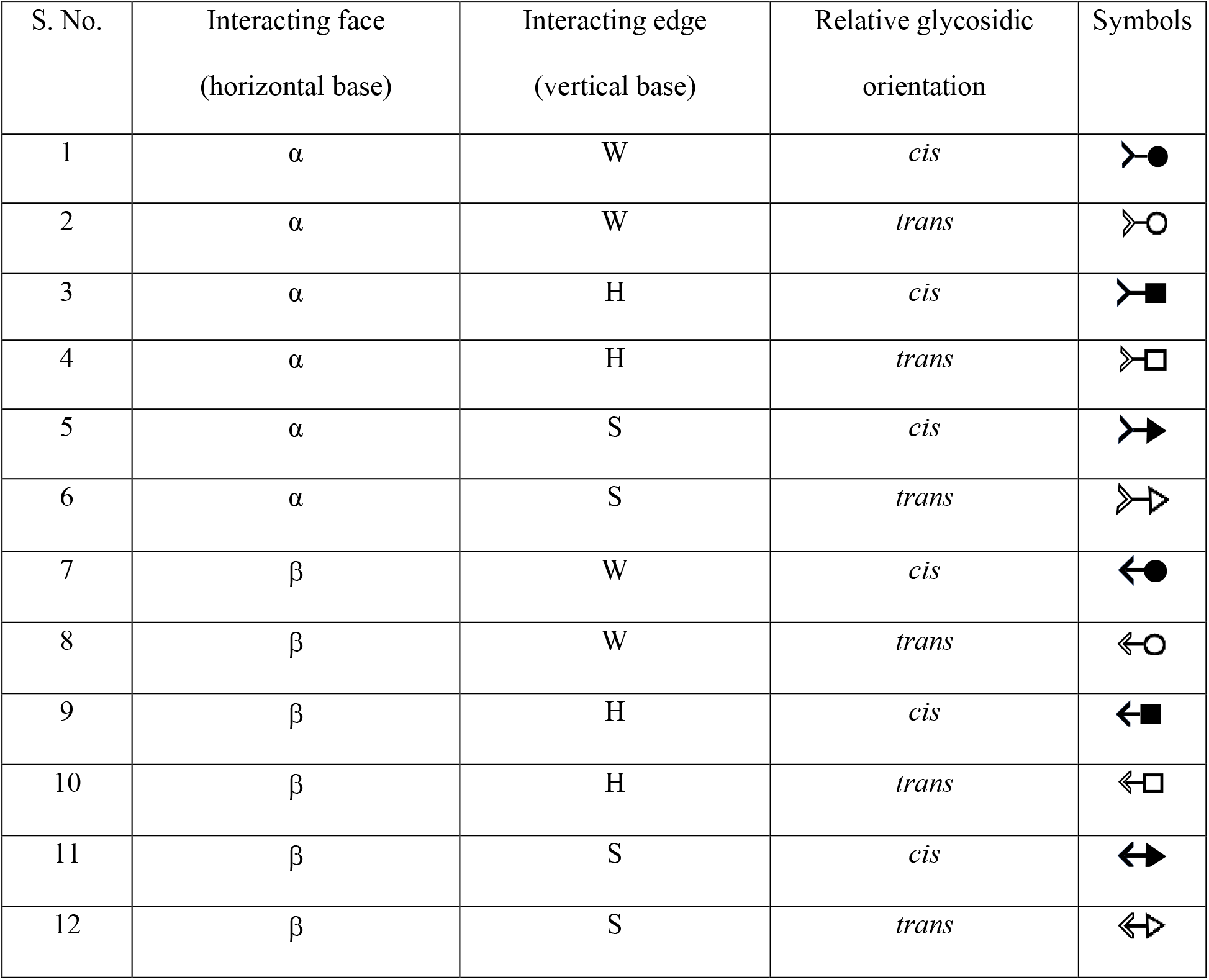
Symbols for representations of 12 geometric families of T-shaped interactions.

| S. No. | Interacting face<br>(horizontal base) | Interacting edge<br>(vertical base) | Relative glycosidic<br>orientation | Symbols |
| --- | --- | --- | --- | --- |
| 1 | $\alpha$ | W | *cis* | |
| 2 | $\alpha$ | W | *trans* | |
| 3 | $\alpha$ | H | *cis* | |
| 4 | $\alpha$ | H | *trans* | |
| 5 | $\alpha$ | S | *cis* | |
| 6 | $\alpha$ | S | *trans* | |
| 7 | $\beta$ | W | *cis* | |
| 8 | $\beta$ | W | *trans* | |
| 9 | $\beta$ | H | *cis* | |
| 10 | $\beta$ | H | *trans* | |
| 11 | $\beta$ | S | *cis* | |
| 12 | $\beta$ | S | *trans* | |

#### Identity of the participating ring of the horizontal purines

Depending on whether the five-membered, six-membered, or both purine rings participate, a T-shaped interaction involving a purine at the horizontal position can be further categorised by adding ‘5’, ‘6’ or ‘56’ to the original face-edge nomenclature (Fig. 2D). For example, the A-C αW *cis* contact involving the interaction of the ‘5’ membered ring of A can be designated as A-C 5 αW *cis*.

Overall, combining six face-edge geometries, three ring identities and two glycosidic bonds, each purine-purine or purine-pyrimidine (purine as the horizontal base and pyrimidine as the vertical base) T-shaped interaction can have 36 (6*3*2) structural possibilities. However, each pyrimidine-purine or pyrimidine-pyrimidine interaction will have 12 (6*2) distinct possibilities. Depending on the base sequence, purine-purine and purine-pyrimidine interactions can have 288 total possibilities (36 each for A-A, A-G, G-A, G-G, A-C, A-U, G-C and G-U), whereas pyrimidine-purine and pyrimidine-pyrimidine interactions can have 96 possibilities in total (12 each for C-A, C-G, U-A, U-G, C-C, C-U, U-C and U-U). This gives rise to a total of 384 distinct theoretically possible T-shaped arrangements (Fig. 4).

**Figure 4.**
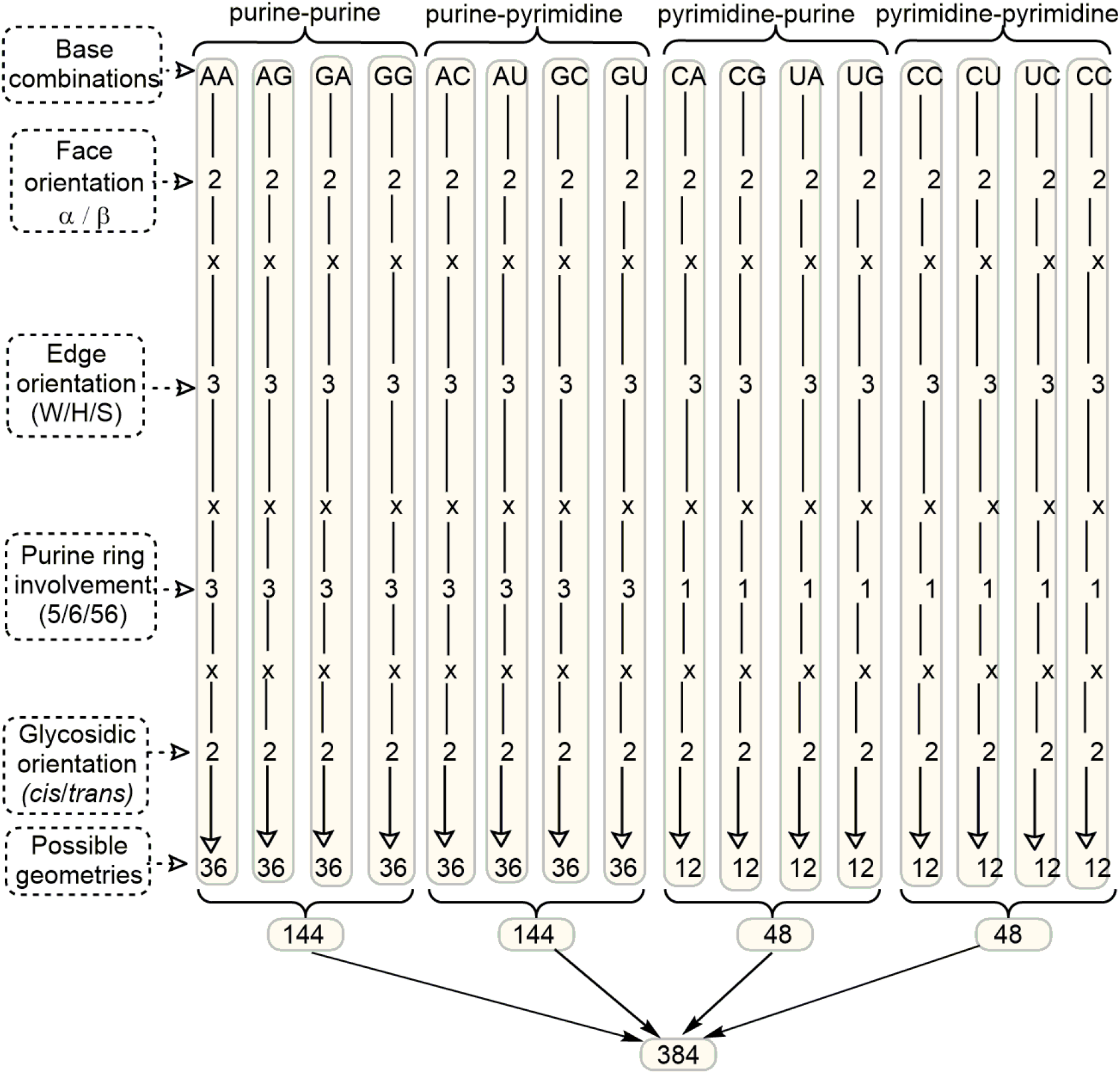
Possible arrangements of the bases in T-shaped interactions, classified with respect to the base-edge orientation, glycosidic orientation and ring involvement of the base providing the face.

Each of these theoretically possible stacking arrangements can occur between nucleotides, which are either in consecutive positions in sequence space or separated by other nucleotides within the sequence space. Thus, depending on the difference between the nucleotide residue numbers, each T-shaped interaction can be annotated as *consecutive, non-consecutive*, or *inter-RNA* (Fig. 2D). Though T-shaped examples between adjacent or non-adjacent nucleotides are not expected to be geometrically different, this final classification can help clarify the functional role these contacts play in different RNA regions.

### Identification of T-shaped interactions in RNA crystal structures

We used the proposed scheme to classify the T-shaped interactions present in the entire data set of RNA crystal structures (see the Materials and Methods section). This open-source python-based identification and classification tool is accessible through github (github.com/PSCPU/T-shaped-interaction). The program generates an output file containing classification of each T-shaped contact present in an RNA PDB file, along with the values of geometrical parameters (i.e., 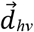, *θ_hν_*, *τ_h_* and *τ_ν_*) that characterize the interaction. A sample output file containing the description of T-shaped contacts identified in a PDB is provided in the Supplementary Information (Supplemental Fig. S1). Consolidated output file containing all identified T-shaped interaction in the total dataset is also provided as a separate supplemental text file.

A total of 3261 instances of T-shaped interactions were found in the dataset of 2286 RNA crystal structures, although these interactions appear in just under one fifth (19.9%) of the structures (Supplemental Table S1). These are not as ubiquitous as the stacking interactions we classified (Jhunjhunwala et al. 2021), so it is suggestive that when present, they might play important roles, especially as they imply a tight 90° kink in an RNA strand. The structures containing T-shaped interactions span all biota, from viruses (e.g., PDB: 2GTT, the nucleoprotein-RNA complex of the rabies virus) to archaea (e.g., 1KQS, the 50S ribosomal subunit of *Haloarcula marismortui*), bacteria (e.g., 1FJG, the 30S ribosomal subunit of *Thermus thermophilus*), and *Homo sapiens* (e.g., 4KR2, the tRNA^gly^-glycyl-tRNA synthetase complex) which points to their evolutionary ubiquity (Fig. 5A). Perhaps unsurprisingly, extant T-shaped interactions are primarily observed in rRNA-derived structures (94.3%), although non-ribosomal structures, mostly involving viral RNA structures (1.3%), tRNA (0.9%), and ribozymes (0.4%), also show T-interactions (Fig. 5B). It is important to note that there is an inherent selection bias in the dataset. Crystallographers are more likely to be interested in specific RNA systems, and certain RNA sequences are also more likely to crystallize. T-structures might be prevalent in other forms of RNA that have not been examined, but with the unusual conformation that they demand, it is reasonable that they would be more common in RNA macromolecules that play a catalytic or complex molecular interaction role; they would not be likely or necessary in most classical coding mRNA for instance.

**Figure 5.**
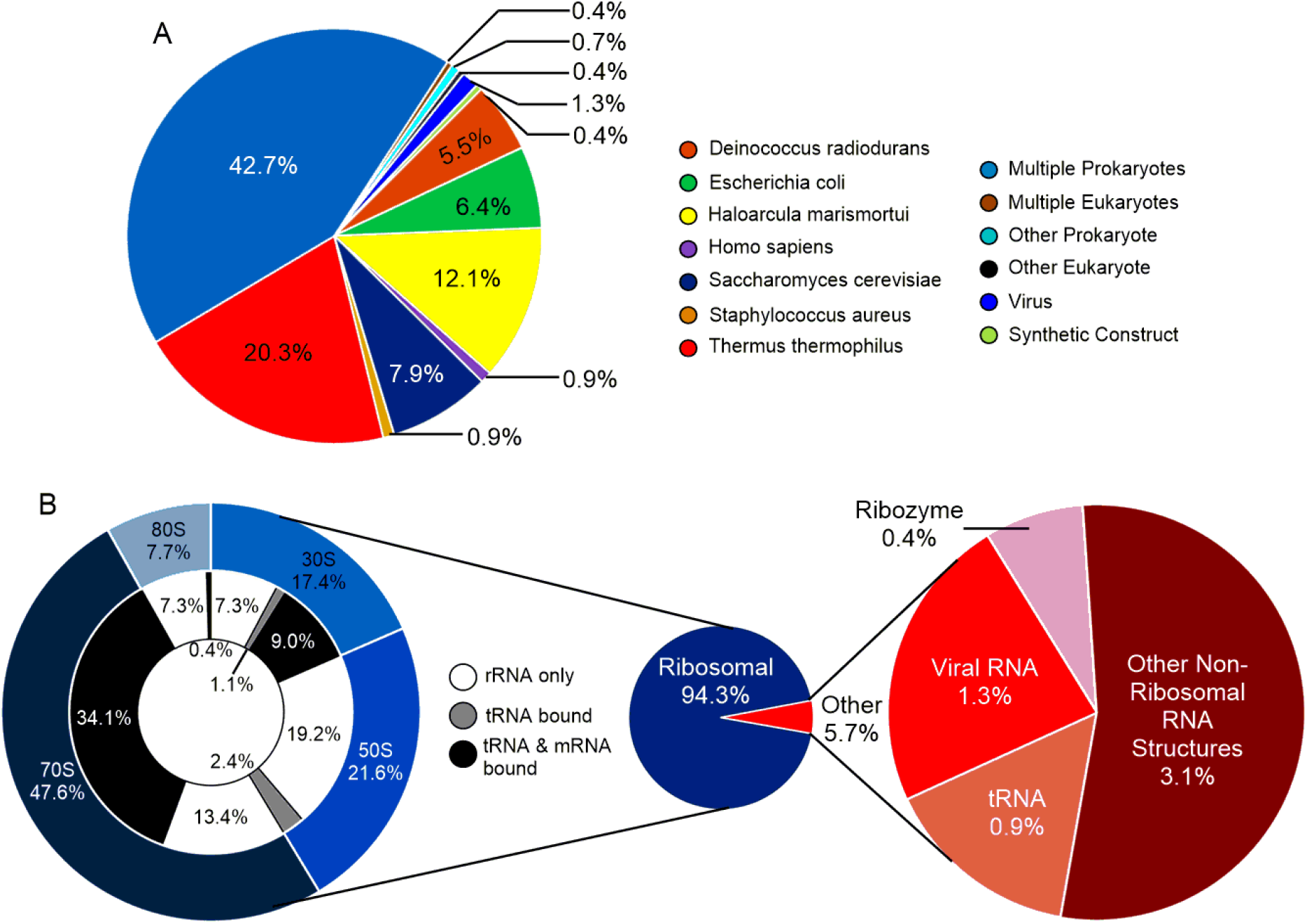
(A) Proportion of PDBs containing T-shaped interactions. (B) Distribution of T-shaped-containing PDBs according to source organism. (C) Distribution of T-shaped-containing PDBs according to RNA type.

In terms of geometrical parameters, the inter-centroid distance 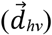 for the majority (71.6%) of T-shaped interactions lies between 4.5 Å and 5.0 Å, with the plurality between 4.8 Å – 4.9 Å. The number of interactions decreases steadily below 4.5 Å (Fig. 6). However, *θ_hν_* remains relatively constant within the interval, with small peaks at 80° to 81° and 88° to 89°, and a small dip in frequency below 78° (Fig. 6B). As expected, all interactions exhibited *τ_a_* values measuring less than 20° and *τ_b_* values measuring over 80° (Fig. 6C). Overall, the narrow distribution of all four geometrical parameters further validates that T-shaped contacts are a distinct phenomenon, and also facilitates the design of an automated search algorithm for the removal of false positives.

**Figure 6.**
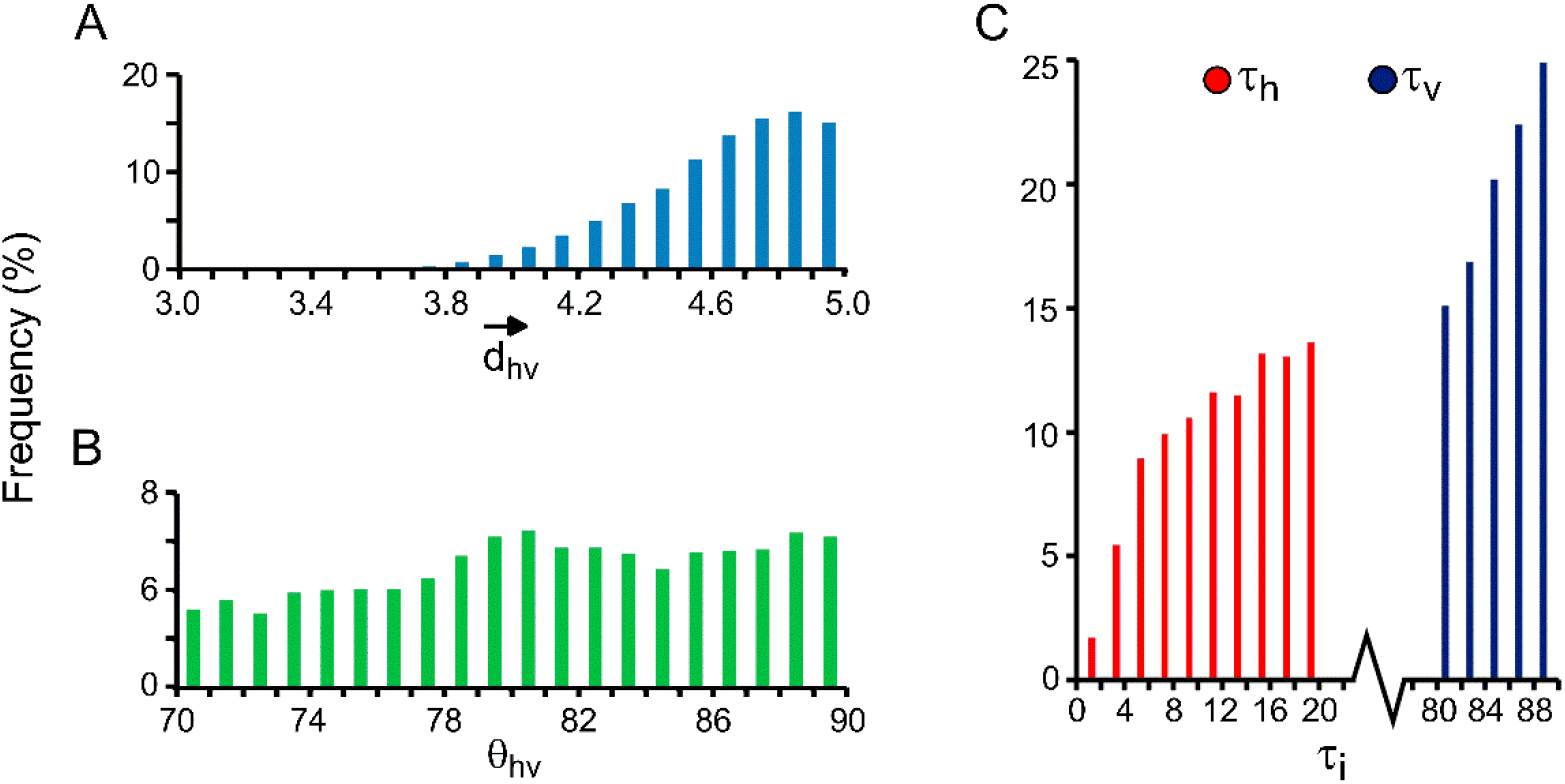
A) Distribution of T-shaped interactions with respect to A) vertical distance (Å) between ring centres, B) tilt angle (θ), and C) angles denoting the relative horizontal displacement of the rings (τ_h_ and τ_v_).

T-shaped interactions slightly prefer the α face (56%) over the β face (44%) of the horizontal base (Supplemental Table S2), and prefer to interact with only one of the two constituent purine rings (6-membered (59%) or 5-membered (40%)) rather than bridging both (Supplemental Table S3). Further, these interactions clearly prefer the H (46%) or S (39%) edge over the W edge of the vertical base (15%, Supplemental Table S4), likely due to the fact that the polar W-edge is generally occupied in classical base pairing interactions (Stombaugh et al. 2009). Alternatively, in terms of the glycosidic orientation, *cis*-oriented interactions are dominant, and comprise around two third of the total interactions (Supplemental Table S5). Finally, T-interactions clearly prefer A as the vertical base (62% of all occurrences), although no specific preference for the horizontal base is observed (Supplemental Table S6).

In terms of sequence combinations, around two-thirds of T-shaped contacts involve two purines (Supplemental Table S2). Further, purine-purine combinations span 85 of the 144 theoretically possible combinations (Fig. 4 and Supplemental Table S7). Included in this purine-purine subset are examples from all 12 possible combinations of the edges (W/H/S), faces (α/β) and glycosidic orientations (cis/trans, Tables 1 and S8). In contrast, only 10% contacts involve a purine at the horizontal position and a pyrimidine at the vertical position (Supplemental Table S2). As a result, these interactions span only 51 of the 144 theoretically possible purine-pyrimidine combinations and only 9 of the 12 possible edges, face, and glycosidic orientations (Tables S8 and S9). An additional 16% contacts involve pyrimidine at the horizontal position and a purine at the vertical positions (Supplemental Table S2). These interactions span 37 of the 48 theoretically possible purine-pyrimidine combinations and involve all 12 possible edge, face, and glycosidic combinations (Tables S8 and S9). Finally, the pyrimidine-pyrimidine interactions contribute only ~9% the total contacts (Supplemental Table S2), although they span 25 of the 48 theoretically possible pyrimidine-pyrimidine combinations and involve 9 of the 12 possible edge, face and glycosidic combinations (Tables S8 and S9). Overall, in terms of the theoretically possible geometrical combinations, crystal structure analysis led to the identification of examples of 198 out 384 possible geometries (Supplemental Table S9 and Fig. 4). Among these, A-G 6 αH *cis* is the most common combination, which along with A-A 5 βS *cis* and G-A 5 αS *trans*, comprises a quarter of all interactions (Supplemental Table S10). It is unclear whether the conformations that are not represented are practically unlikely, or in some cases essentially impossible; the dataset is simply too small to be expected to encompass all possibilities, and this exercise could be repeated after a suitable interval to determine whether these missing conformations are observed as more structures are deposited.

Around two-thirds of the T-shaped interactions involve *non-consecutive* nucleotides (Supplemental Table S11). Over 40% of these *non-consecutive* interactions are separated by only 1 or 2 nucleotides in the sequence space, whereas only 3.5% are separated by 3 to 10 nucleotides. The remaining interactions are separated by 11 to 2150 nucleotides, with noticeable clusters between *n*+*10* and *n*+*20*, *n*+*120* and *n*+*150*, as well as at *n*+*380* (Fig. 7). However, despite their likely extremely important functional role (*vide infra)*, the frequency of *inter-RNA* T-shaped interactions is statistically insignificant.

**Figure 7.**
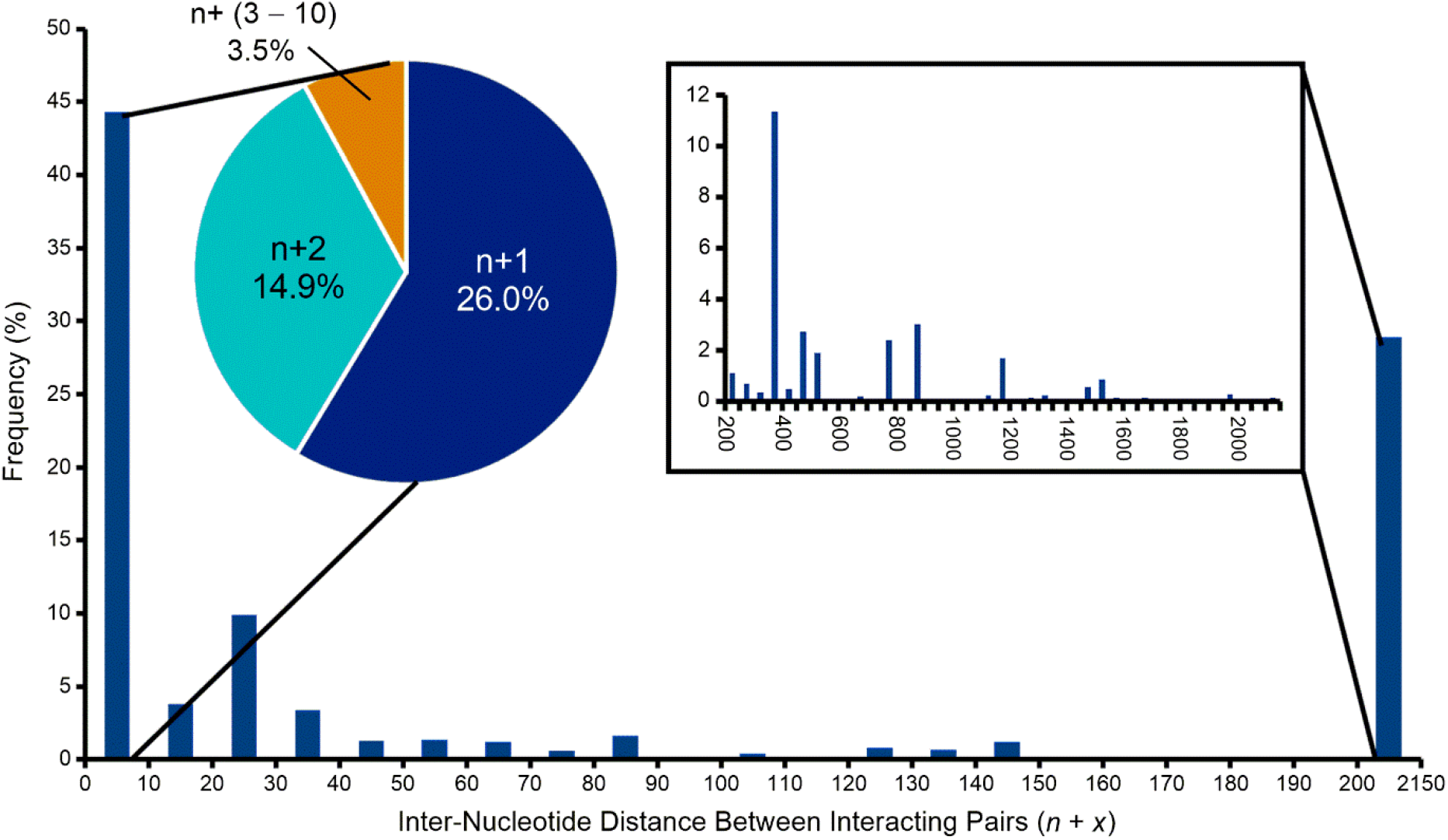
Inter-nucleotide distance between *non-consecutive* T-shaped interacting pairs. Distances from *n* + 1 to *n* + 10 are detailed in the pie chart and large distances from *n* + 200 to *n* + 2150 in the small histogram.

#### Comparison with parallel stacking

For comparison, the dataset was also searched for the analogous parallel face-to-face stacking contacts using the previously developed analogous detection methodology (Jhunjhunwala et al. 2021). Owing to their general role in stabilizing RNA structures, parallel stacking interactions occurs in all RNA-containing PDB files. However, 80% of the PDB structures that involve stacking interactions do not have T-shaped interactions. Overall, the occurrence of stacking interactions is 568 times greater than T-shaped interactions (Supplemental Table S1). This correlates with the evidence that T-shaped contacts occur primarily in specific RNA regions with irregular backbone topologies (*e.g*., loops and junctions).

Further comparison of the statistical distribution of stacking and T-shaped interactions reveals salient differences. Specifically, in terms of the preference of the constituent nucleobases, stacking interactions show reduced preference towards purine-purine combinations (45.5%) compared to T-shaped interactions (65%, Supplemental Table S12). Further, the contribution of pyrimidine-pyrimidine combinations in T-shaped interactions (8.8%) is significantly lower than stacking contacts (20.7%, Supplemental Table S12). However, in terms of the identities of the constituent bases forming the stack, the A-G combination dominates both stacking and T-shaped contacts (Supplemental Table S13). Alternatively, in terms of the glycosidic orientation, although T-shaped interactions involve significant *trans* orientation (33.2%,), stacking interactions are overwhelmingly *cis* oriented (91.6%, Supplemental Table S14). Further, although T-shaped interactions predominantly occur between *non-consecutive* nucleotides (66%), only a quarter of the stacking interactions are *non-consecutive* (Supplemental Table S15). However, due to their rarity in the structure space, *inter-RNA* contacts contribute insignificantly to the overall statistics of both stacking and T-shaped interactions. Overall, this comparison reveals that the physicochemical and structural requirements for a T-shaped interaction are different than those needed to establish parallel stacking interactions.

### Annotation of T-shaped Contacts in RNA Structures

As previously suggested for base pairs (Leontis and Westhof 2001) and stacking interactions (Jhunjhunwala et al. 2021), the annotation of T-shaped interactions in pictorial representations of RNA structures or substructures would help better visually recognize the noncovalent interactions in RNA motifs. In this context, we propose a symbolic formalism for the annotation of T-shaped interactions that combines the features of Leontis and Westhof’s base-pairing annotations (Leontis and Westhof 2001) and Jhunjhunwala *et al’s* annotations of parallel stacking interactions (Jhunjhunwala et al. 2021). Specifically, we propose that the edge of the vertical base should be specified using the symbols used for specifying the edges of nucleobases in pairing interactions (*i.e*., circle for the W edge, square for the H edge and triangle for the S edge). However, the interacting face of the vertical base can be designated using an arrowhead (› for the α face and   for the β face), which can be joined by a dash (–) to complete the assignment of a T-shaped interaction. Further, these symbols can be used in their filled and hollow forms to specify *cis* and *trans* glycosidic orientations, respectively. For instance, both symbols 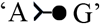 and 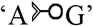 would represent T-shaped interaction involving the interaction of horizontal A through its α-face and vertical G through its W-edge, albeit with *cis* and *trans* glycosidic orientations, respectively. Symbols for all the 12 families of T-shaped interactions are provided in Table 1.

When a horizontal purine is involved in a T-shaped interaction, information specified by the above symbols can be further enriched by providing information on the identity of the constituent purine ring (‘5’, ‘6’ or ‘56’). To illustrate how these symbols can denote T-shaped interactions, we present one A-A example from each of the 12 geometric families (Fig. 8).

**Figure 8.**
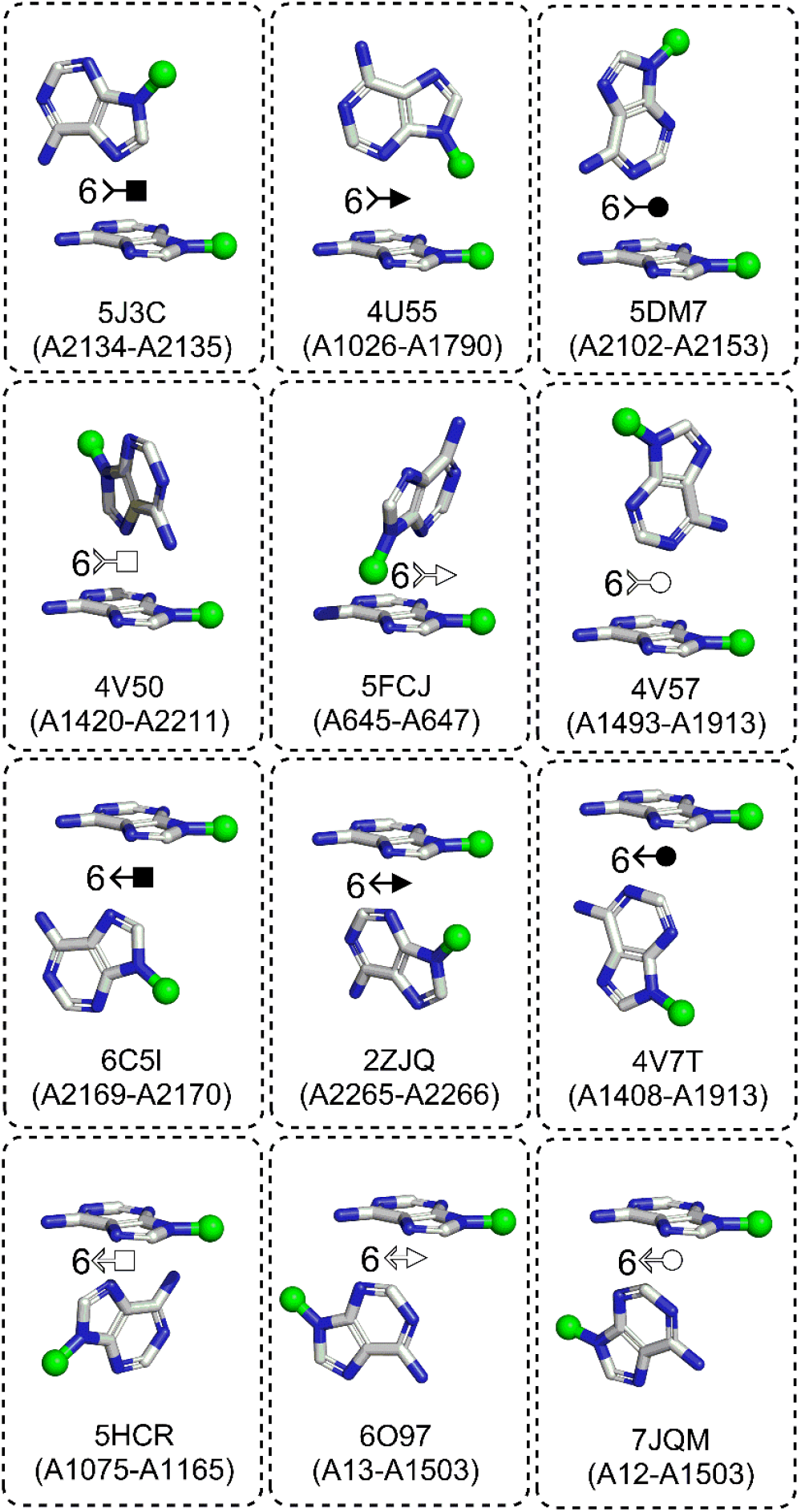
Illustration of the symbolic representation of edge-face nomenclature for T shaped interactions, using one example from each of the 12 families of adenine-adenine combinations (Table 1). PDB code and interaction code (horizontal base-vertical base) is provided for each interaction.

### Structural context of occurrence of selected T-shaped interactions in RNA structures

To aid in visual understanding of how the proposed classification scheme can be used to enrich our understanding of T-shaped interactions in RNA 3D structures, we annotate these interactions in certain specific RNA motifs (Fig. 9). Specifically, we used one example structure from each of the major RNA classes in which these interactions were identified (i.e., tRNA, viral RNA, ribozyme and ribosome). First, we annotate the βH *trans* (*i.e*., 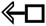) T-shaped interaction between U60 and A58 belonging to the T loop, which likely plays a role in structuring this loop of tRNA^Gly^ in its crystal structure in complex with the glycyl tRNA synthetase (PDB code: *4KR2*, Fig. 9A). A sharp bend in the strand leads to the *‘trans”* arrangement of the interacting bases, which contrasts the ‘*cis*’ orientation of the βS arrangement (*i.e*., 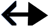) of the intraloop C11-A14 of the vitamin B12 mRNA aptamer (*1DDY*, Fig. 9B). Interestingly, a similar ‘*trans*’ arrangement is observed in the 5 αH A231-A259 (*i.e*., 5 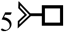) interaction present in the three-way junction of the lariat capping ribozyme (*4P95*, Fig. 9C). However, the *‘consecutive”* A70-A69 5αW interaction present in the circular viral RNA strand of the rabies virus nucleoprotein-RNA complex acquires a *“cis”* 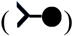 arrangement (*2GTT*, Fig. 9D). Similarly, the *“consecutive”* G1033-G1032 6 βH and the *‘non-consecutive”* U82-U85 βH intraloop interactions, as well as the A975-A1357 5 βS loop-loop interactions with the crystal structure of 70S ribozyme of *Thermus Thermophilus* acquire the *“cis”* arrangements (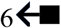 and 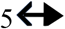 respectively, *6BOH* Fig. 9E).

**Figure 9.**
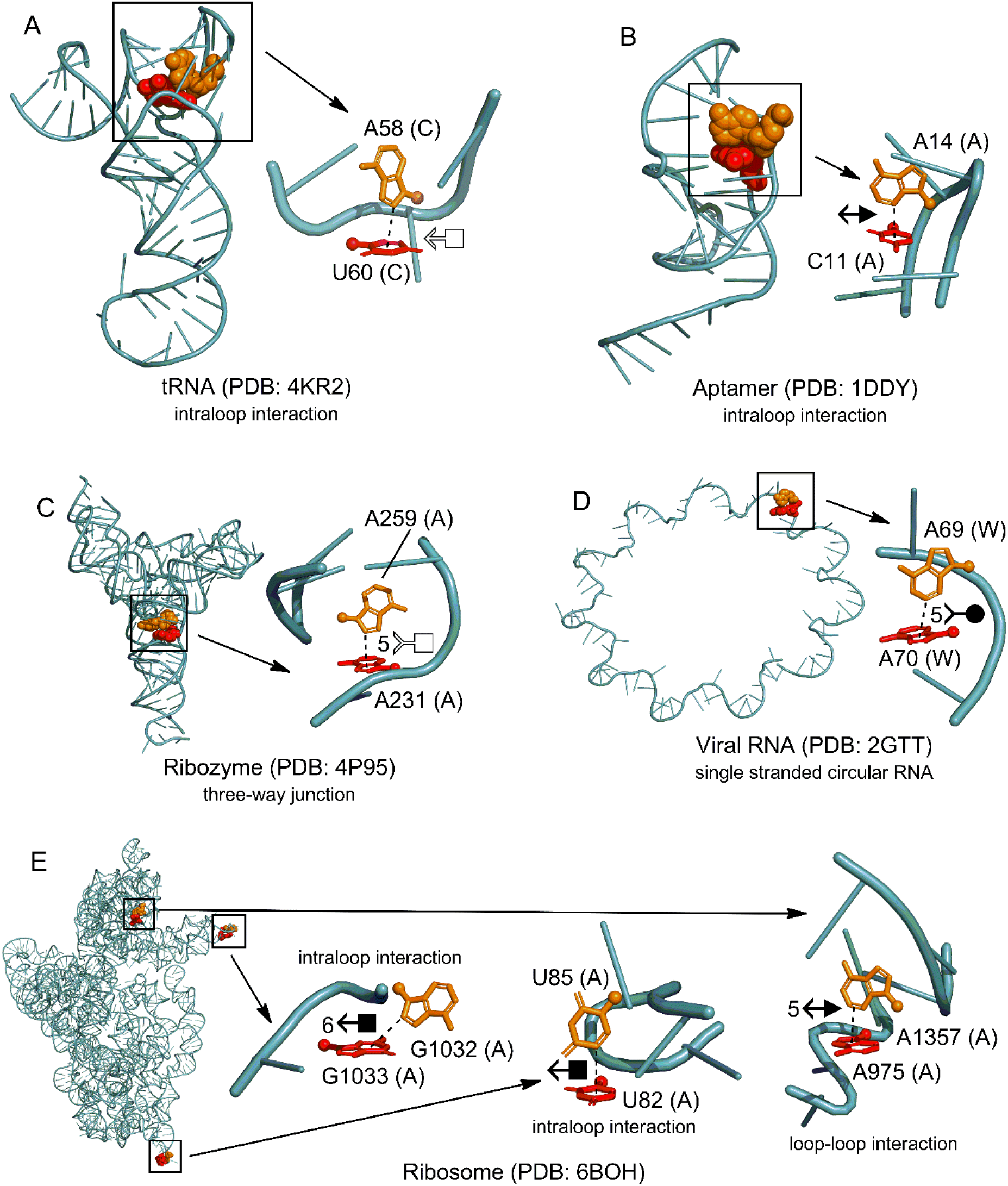
Sample T-shaped interactions from various PDBs with indicated nucleotide type, residue number, and chain number, respectively.

In addition to T-shaped contacts between bases belonging to the same RNA sequence, out study identified a unique T-shaped contact between two different RNA elements in the crystal structure of the hepatitis A virus IRES domain V (dV) in complex with a synthetic antibody (Fig. 10, *6MWN)*. The βH *cis* geometry of this U626(A)-U626(B) T-shaped interaction is like the G1033-G1032 and U82-U85 geometries in the ribosome (Fig. 9E). Together with intra-RNA T-shaped contacts, the existence of an *inter-RNA* T-shaped contact suggests that these interactions may play a role in noncovalently structural organization within RNA, as well as between two RNA structures.

**Figure 10.**
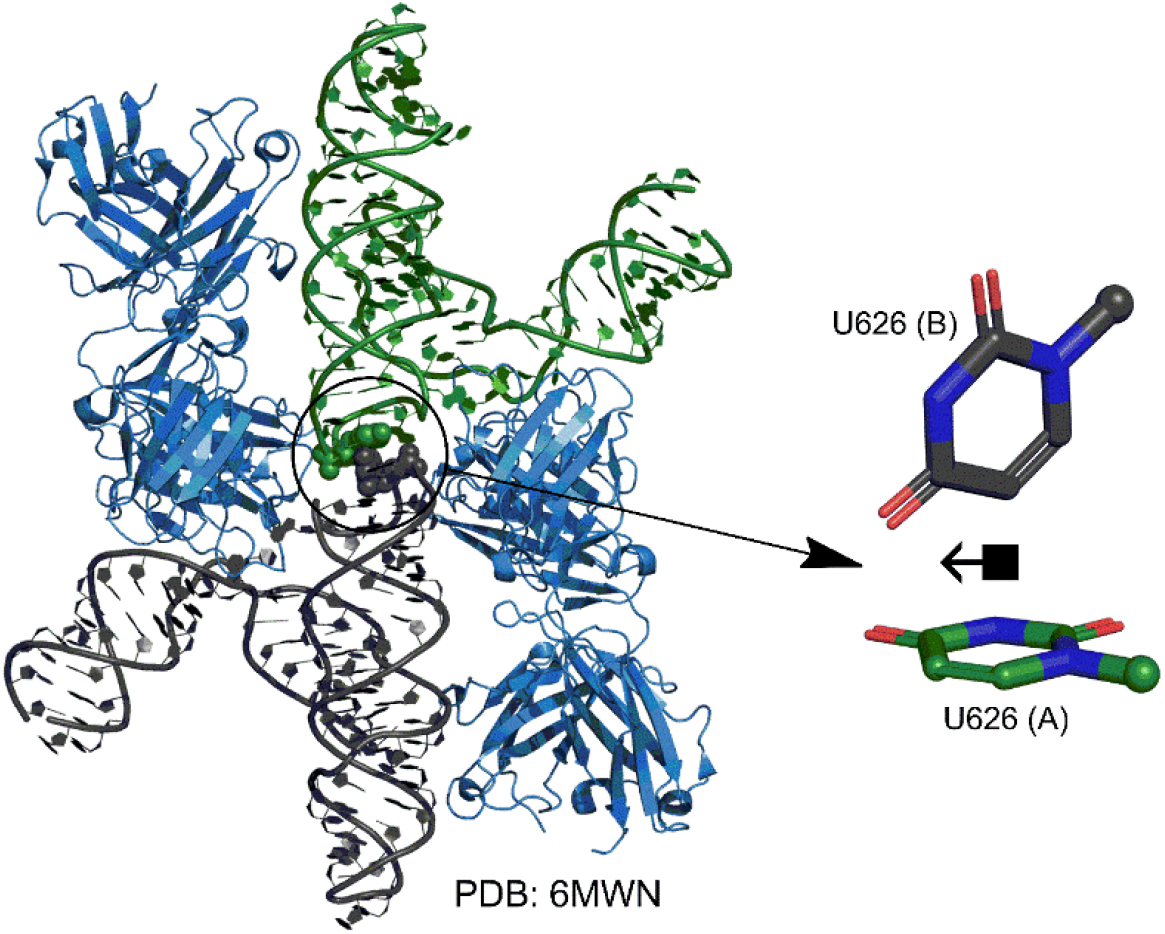
Representative structure of a crystal structure *inter-chain* RNA T shaped interaction.

## CONCLUSION

In the present work, we developed an automated tool for the identification of hitherto unknown T-shaped contacts that occur between nucleobases in RNA structures. We further developed and applied a comprehensive classification scheme to unambiguously categorize these contacts in RNA structures. In addition, we proposed a symbolic representation to annotate T-shaped interactions in secondary structure diagrams and 3D structural representations of RNA. Thus, our study adds to the list of previously-established noncovalent effects, including but not limited to, base pairing, parallel base stacking, lone pair-π interactions between ribose and nucleobases, lone pair-π and OH–π interactions between water and nucleobases, base-wedged and base intercalated stacking, as well as base-phosphate interactions, that help organize RNA structures. This effort, by providing a language and a tool to discuss these interactions can help with identifying and understanding their role in RNA organization.

## MATERIALS AND METHODS

### Structural dataset

A data set containing all RNA crystal structures deposited in the Protein Data Bank (PDB) through May 28, 2022, was used to identify the T-shaped interactions. The structures were searched from the PDB by setting the ‘Polymer Entity Type’ to ‘RNA’and by choosing ‘X-ray’ in the “Method” tab. Further, in agreement with previous studies, the refinement resolution of the crystal structures was set to ≤ 3.5 Å (Chawla et al. 2014; Wilson et al. 2014; Chawla et al. 2015). A total of 3280 structures were obtained from this search, which were futher filtered using the CD-HIT suite (Huang et al. 2010), to retain only a single representative from clusters of similar structures with >80% similarity. This procedure lead to the final 2286 crystal structures (Supplementary Supplemental Table S16).

### Identification of T-shaped interactions

In analogy with the ‘Stackdetect’ algorithim previously developed for detecting base-base stacking interactions (Jhunjhunwala et al. 2021), a python script was developed for automated detection and classification of T-shaped interactions in RNA crystal structures. In the first step, the Cartesian coordinates of atoms forming the five-membered ring skeleton of a purine (*i.e*., C4, C5, N7, C8 and N9 atoms) and the six-membered ring of a purine or a pyrimidine (N1, C2, N3, C4, C5 and C6), and C1’ of the sugar moity were extracted, and the coordinates of the centroid of each ring was determined (*x_c_, y_c_*, z_c_).

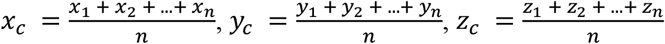

Here (*x*_1_, *y*_1_, *z*_1_), (*x*_2_, *y*_2_, *z*_2_), (*x*_3_, *y*_3_, *z*_3_)…… (*x_n_, y_n_, z_n_*) are the cartesian coordinates of atoms of each 5-membered purine ring or 6-membered purine or pyrimidine ring. The centroid coordinates were then used to determine the position vector 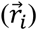 of each (*i^th^*) ring atom from the ring centroid. Specifically:

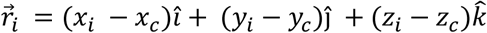

Here (*x_i_*, *y_i_*, *z_i_*) denote the cartesian coordinates of the atom *i*. Subsequently, the distance vector 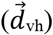 that connects the centroids of vertical ring (denoted as ring ‘v’) and the horizontal ring ((denoted as ring ‘h’)) that participate in a T-shaped interaction was determined using the centroid coordinates of the rings ((*x_hc_, y_hc_, z_hc_*) and (*x_νc_*, *y_νc_*, *z_νc_*), Fig. 11A). Specifically:

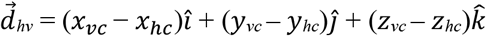

**Figure 11.**
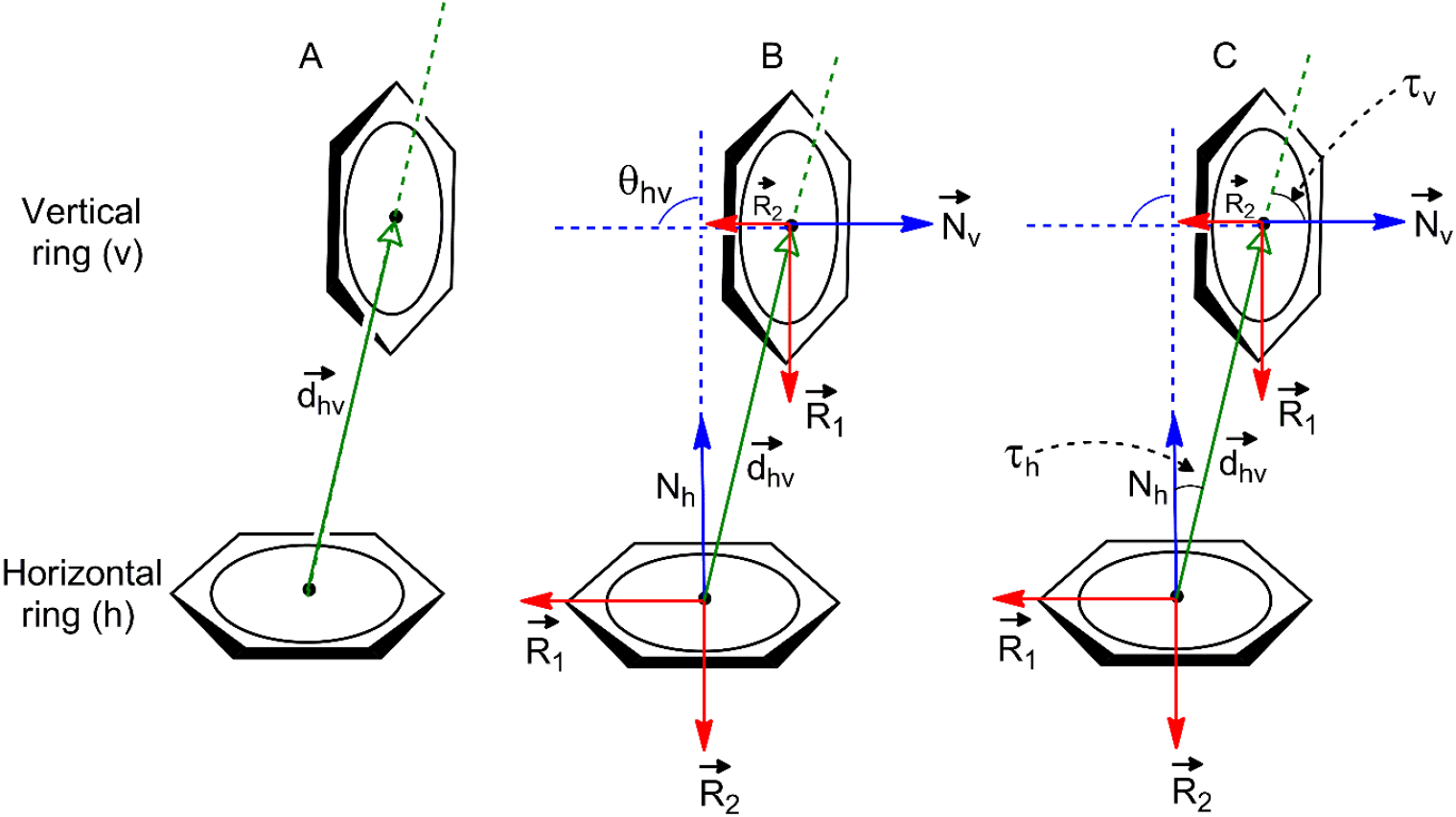
Representation of geometrical parameters used for locating T-shaped interactions in RNA crystal structures.

For each interacting ring, a mean plane was then defined using the two orthogonal vectors 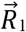 and 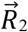 using the Cremer and Pople method (Cremer and Pople 1975).

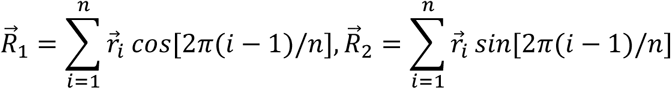

The vectors normal to the mean plane vectors (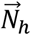 and 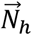) of each ring were then determined as the cross product of 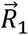 and 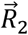.

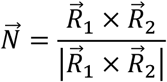

The inverse cosine of the dot product of the normals 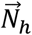 and 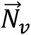 of the interacting rings *h* and *ν* was used to calculate *θ_hν_*, the tilt angle between the rings (Fig. 11B (Cremer and Pople 1975)). Specifically:

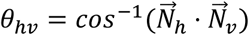

Further, the angle between 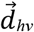 and 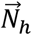or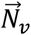(i.e., *τ_h_* or *τ_ν_*, Fig. 11C) was calculated for each interaction.

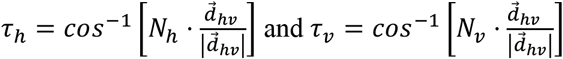

The values of *τ_h_* and *τ_ν_* determine the relative horizontal displacement of the interacting rings.

In line with the previous studies on T-shaped interactions between DNA nucleobases and aromatic amino acids (Wilson et al. 2014), following criteria was used for locating a T-shaped interaction between the two interacting rings:

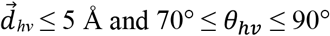

However, the ranges for *τ_h_* and *τ_ν_* needed to be determined empirically through preliminary trials. The best cut offs for these interactions to retain a T-shaped conformation were found to be:

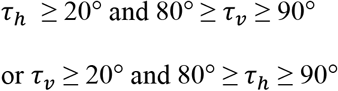

For purines, these parameters were calculated separately for both 5-membered and 6-membered rings, and the identity of the interacting ring was assigned accordingly (*vide infra)*.

### Classification of T-shaped interactions

For each T-shaped interaction, the vertical base can interact with the horizontal base through one of its three edges – the Watson-Crick (W) edge, Hoogsteen (H) edge or the Sugar (S) edge (Fig. 2A (Leontis and Westhof 2001)). The interacting edge of the vertical base was computationally identified by calculating the coordinates of the centre of each edge of the base by averaging the coordinates of all donor and acceptor atoms that constitute the edge (Supplemental Fig. S2 and Supplemental Table S17 (Leontis and Westhof 2001)).

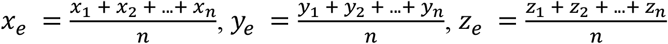

Here, (*x*_1_, *y*_1_, *z*_1_), (*x*_2_, *y*_2_, *z*_2_), (*x*_3_, *y*_3_, *z*_3_)…… (*x_n_, y_n_, z_n_*)) are the Cartesian coordinates of the selected edge atoms and *x_e_*, *y_e_, z_e_* are the coordinates of the edge centre (Supplemental Table S17). Subsequently, the distance between the centroid of the horizontal ring and the centre of each edge of the vertical base, was calculated. The edge closest to the centroid of the interacting face of the horizontal ring was accordingly assigned as the “interacting edge” for the purposes of classifying the T-shaped interaction.

The identity of the interacting face of the horizontal ring was then established. Specifically, the face was designated as ‘α’ when the standard nucleobase atomic numbering of the ring from lowest numbered atom to the subsequent higher numbered atoms by shortest route progresses in a ‘clockwise’ manner, when the base is viewed from the top of the interacting face (Fig. 2B (Rose et al. 1980)). Alternatively, if the progression occurs in a ‘counter clockwise’ direction, the face is named as ‘β’ (Fig. 2B). However, by convention, the face designation of a purine is derived from the numbering of its six membered ring, irrespective of whether the six-membered or the five-membered ring interacts with the other base (Rose et al. 1980). Computationally, the face assignment is done using the direction of the normal to the mean plane vector of the horizontal base 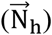. As the mean plane is defined by the positions of ring atoms (N1 to N6, Fig. 11), the direction of the normal vector 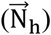 is such that it always extends from the β face (Rose et al. 1980). For this reason, the face is designated as α if the angle between 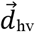 and 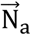 is greater than 90°, otherwise it is assigned as β.

The mutual glycosidic orientation of each T-shaped interaction was assigned by calculating the angle *σ_hν_* (∠(C1’)_*h*_–X_*h*_–X_*ν*_–(C1’)_*ν*_), where (C1’)_*h*_ and (C1’)_*ν*_ represent the C1’ atoms of the nucleotides that include the horizontal and the vertical ring, respectively, and X_*h*_ and X_*ν*_ are the centroids of the respective base rings that connect to the sugar through the glycosidic bond (5-membered ring in case of purines and 6-membered ring in case of pyrimidines, Fig. 2C).

The identity of the interacting ring of the horizontal base was determined by calculating the distance between the centroid of the vertical base and each ring (5- or 6-membered ring of purines and 6-membered ring of pyrimidines) of the horizontal base. For example, if the geometrical parameters (i.e., 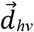, *θ_hν_*, *τ_h_* and *τ_ν_*) of the centre of the vertical base and the centre of 5-membered ring of the horizontal purine fall under the cut-off value, the interaction is denoted as ‘5’ (Fig. 2D). Alternatively, if the parameters of the 6-membered purine ring falls within the cut-off, the ring topology is assigned as ‘6’ (Fig. 2D). However, if the parameters for both 5-memebered and 6-membered purine rings fall with the cut off, the T-shaped interaction is noted as ‘56’ (Fig. 2D).

Each T-shaped interaction was further annotated as *consecutive*, *non-consecutive*, or *inter-RNA*, based on the absolute value of the difference between the nucleotide residue numbers in the PDB file, and whether the interacting nucleotides belong to the same RNA entity or to two different interacting RNA strands. A T-shaped interaction between nucleotides belonging to the same chain in the PDB file is annotated as *consecutive*, if the difference between their residue numbers equals 1, i.e., they are immediately adjacent; otherwise, the interaction is annotated as *non-consecutive*. However, if the interacting nucleotides belong to two different RNA present in the same PDB file, the interaction is classified as *inter-RNA* (Fig. 2E).

## SUPPLEMENTAL MATERIAL

Supplemental material is available for this article.

## ACKNOWLEDGMENTS

The research project was supported by Department of Science and Technology (DST), University Grants Commission (UGC). P.S. thanks the Department of Science and Technology (DST), and University Grants Commission (UGC), New Delhi, for financial support through the DST INSPIRE (IFA14-CH162) and the UGC FRP (F.4-5(176-FRP/2015(BSR)) programs, respectively. JFT thanks the Natural Sciences and Engineering Research Council of Canada. Z.A. thanks UGC for a Senior Research Fellowship.

